# Interleukin-3 is a predictive marker for severity and outcome during SARS-CoV-2 infections

**DOI:** 10.1101/2020.07.02.184093

**Authors:** Alan Bénard, Anne Jacobsen, Maximilian Brunner, Christian Krautz, Bettina Klösch, Izabela Swierzy, Elisabeth Naschberger, Torsten Birkholz, Ixchel Castellanos, Denis Trufa, Horia Sirbu, Marcel Vetter, Andreas E. Kremer, Kai Hildner, Andreas Hecker, Fabian Edinger, Matthias Tenbusch, Enrico Richter, Hendrik Streeck, Marc M. Berger, Thorsten Brenner, Markus A. Weigand, Filip K. Swirski, Georg Schett, Robert Grützmann, Georg F. Weber

**Author notes:** Correspondence to: Georg F. Weber, Or Alan Bénard, Department of Surgery, Universitätsklinikum Erlangen, Friedrich-Alexander University (FAU) Erlangen-Nürnberg Erlangen, Germany.

## Abstract

Severe acute respiratory syndrome coronavirus 2 (SARS-CoV-2) is a worldwide health threat. Here, we report that low plasma interleukin-3 (IL-3) levels were associated with increased severity and mortality during SARS-CoV-2 infections. IL-3 promoted the recruitment of antiviral circulating plasmacytoid dendritic cells (pDCs) into the airways by stimulating CXCL12 secretion from pulmonary CD123^+^ epithelial cells. This study identifies IL-3 as a predictive disease marker and potential therapeutic target for SARS-CoV-2 infections.

## Main Text

Coronavirus Disease 2019 (COVID-19) is an acute infection of the respiratory tract that had spread worldwide^1^. As patients may develop severe respiratory syndrome^2^, it is important to define predictive markers allowing clinicians to identify patients at risk at an early stage of the disease. IL-3 has been described as an important immune mediator during infections^3,4^. We therefore investigated whether IL-3 might influence the outcome of SARS-CoV-2 infection.

Patients with severe COVID-19, characterized by necessity for intensive care treatment and high plasma CRP levels (Extended Data Fig. 1a), had reduced plasma IL-3 levels as compared to patients with non-severe COVID-19 or patients that had recovered (Fig. 1a). As well, patients with high viral load presented lower plasma IL-3 levels as compared to patients with low viral load (Fig. 1b), both results suggesting an association between plasma IL-3 levels and disease severity in SARS-CoV-2 infections. In this prospective multicentric observation study, Kaplan-Meier survival analysis revealed that plasma IL-3 levels may predict the outcome of SARS-CoV-2 infections: using a minimal p-value approach, patients with plasma IL-3 levels <20 pg/ml at admission had a poorer prognosis as compared to patients with plasma IL-3 levels ≥20 pg/ml at admission (Fig. 1c and Extended Data Table 1), this association remaining significant after adjusting for prognostic parameters in multivariate analysis (Extended Data Table 2). Older age was described to be associated with greater risk to develop severe COVID-19^5^. Patients older than 65 years showed reduced plasma IL-3 levels as compared to patients younger than 65 years (Fig. 1d). Thus, the analysis of plasma IL-3 levels and patient age allowed to identify three groups at risk to die from COVID-19: patients <65 years with plasma IL-3 levels ≥20 or <20 pg/ml had a low to intermediate risk to die whereas patients ≥65 years with plasma IL-3 levels <20 pg/ml had a high risk to die (95%-CI: 1.680 – 118.218; OR: 14.091) (Fig. 1e, Extended Data Fig. 1b and Extended Data Table 3). Survivors from severe SARS-CoV-2 infection displayed higher circulating pDC numbers over time as compared to non-survivors whereas no differences could be detected in the numbers of circulating neutrophils (Fig. 1f). SARS-CoV-2^+^ patients exhibiting either high plasma IL-3 levels or high circulating pDC numbers had therefore a better prognosis, suggesting a putative link between IL-3 and pDCs during infection. Bronchoalveolar lavage fluid (BALF) analysis (Extended Data Table 4) revealed that patients with high levels of IL-3 showed increased percentages of pDCs as compared to those with low levels (Fig. 1g).

**Figure 1.**
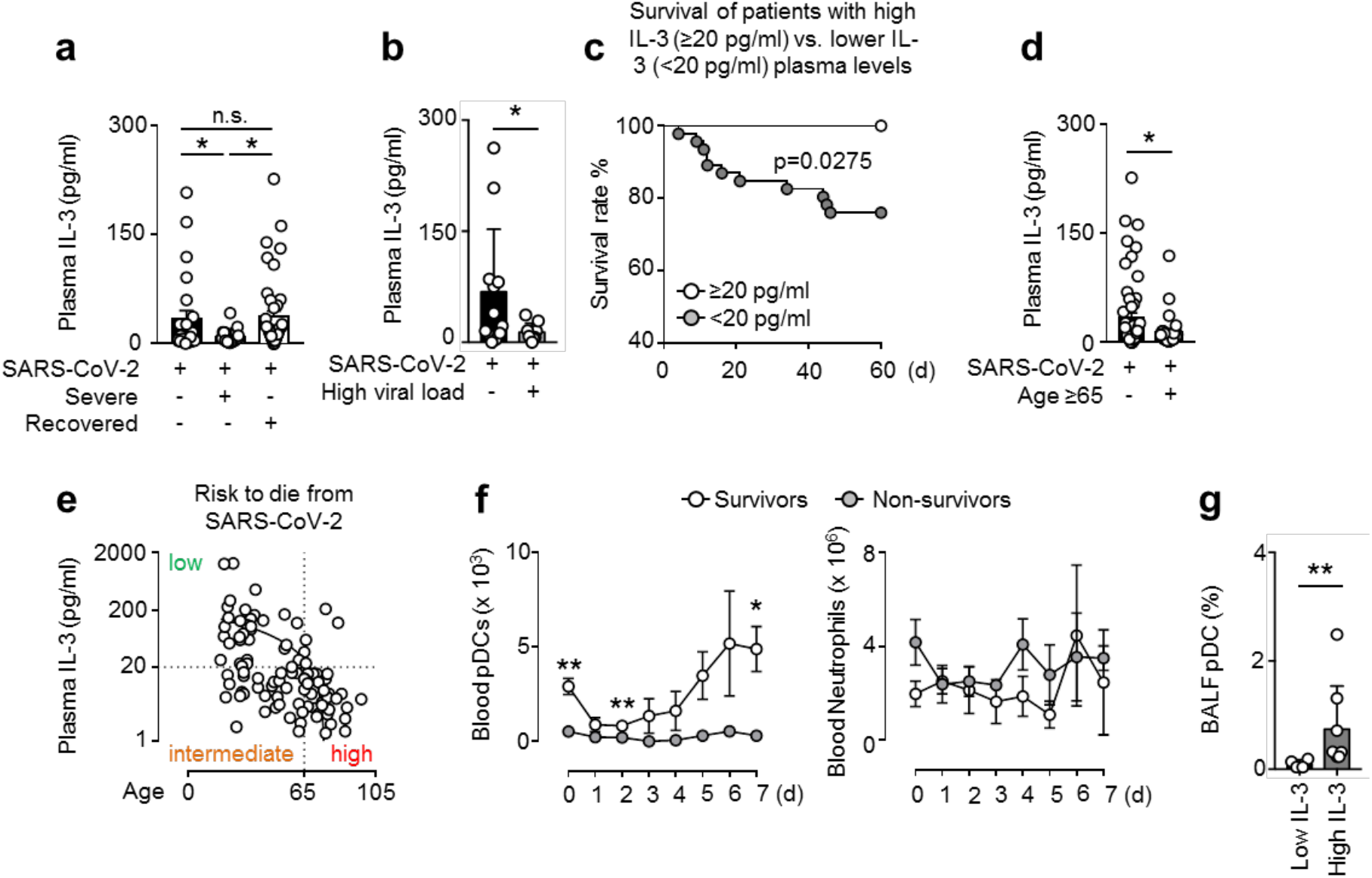
Low plasma interleukin-3 levels are associated with severity and outcome in COVID-19. **a,** Plasma IL-3 levels of SARS-CoV-2^+^ patients with or without severe disease and in patients that had recovered from infection. One-way ANOVA. n=106. **b**, Plasma IL-3 levels of SARS-CoV-2^+^ patients with or without high viral load. n=21. **c**, Kaplan-Meier analysis showing the survival of SARS-CoV-2^+^ patients with high (≥20pg/ml) or low (<20pg/ml) plasma IL-3 levels (measured within 24 hrs after admission). n=64. **d**, Plasma IL-3 levels of SARS-CoV-2^+^ patients older or younger than 65 years. n=106. **e**, Correlation between plasma IL-3 levels and age. n=106. **f**, Absolute numbers of circulating pDCs and neutrophils in SARS-CoV-2^+^ patients from their admission to ICU and 1 to 7 days later. n=9. **g**, Percentage of pDCs in BALF of patients with pulmonary diseases with high or low BALF IL-3 levels. n=13. Data are mean ± s.e.m., *P < 0.05, **P < 0.01, ***P < 0.001, unpaired, 2-tailed Student’s t test using Welch’s correction for unequal variances was used.

We next investigated the link between IL-3 and the amount of pDCs in the airways experimentally. Upon intranasal (i.n.) IL-3 administration, naive WT mice displayed higher absolute numbers of pDCs, but not neutrophils, in the lung parenchyma as compared to controls (Fig. 2a and Extended Data Fig. 2a) as well as substantially higher levels of IFNα and IFNλ in the BALF after subsequent i.n. CpG injection (Fig. 2b). Depletion of pDCs induced a strong reduction of IFNα levels in BALF of mice pre-treated with IL-3 upon i.n. CpG administration, whereas IFNλ secretion was only partially impaired (Extended Data Fig. 2b-c). Among the chemokines known to drive pDC migration into inflamed tissues^6^, only *Cxcl12* expression was increased by IL-3 treatment in lungs of mice upon CpG administration (Fig. 2c and Extended Data Fig. 3a). We also detected increased CXCL12 levels in the BALF of WT mice treated with IL-3 (Fig. 2d) and in the supernatant of *ex vivo* cultured lung cells derived from naive WT mice upon IL-3 stimulation (Extended Data Fig. 3b). The induction of CXCL12 was mediated through the IL-3 receptor common β-chain (CD131), as no increase in CXCL12 levels was observed in the BALF of *Cd131*^*−/−*^ mice upon IL-3 stimulation (Fig. 2d).

**Figure 2.**
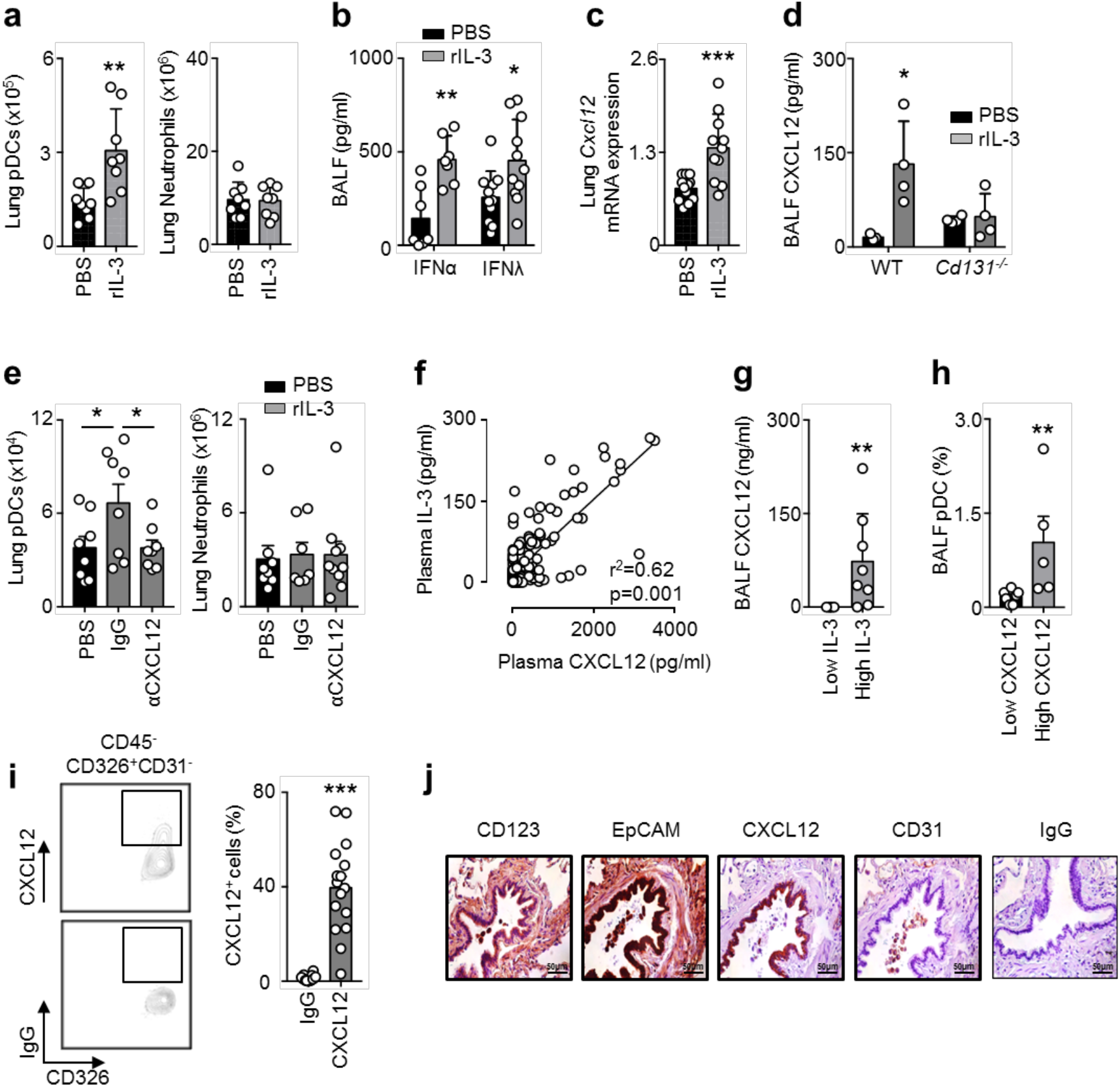
Interleukin-3 promotes the recruitment of pDCs into the lung in a CXCL12-dependent manner. **a**, Absolute numbers of pDCs and neutrophils in the lungs of naive mice 24 h after the i.n. injection of PBS or IL-3. n=8. **b**, Levels of IFNα and IFNλ in the BALF of naive mice that received an i.n. injection of PBS or IL-3, followed by an i.n. injection of CpG 8 h later, and were sacrificed 16 h later. n=7-11. **c**, Relative mRNA expression of *Cxcl12* in the lungs of naive mice 24 h after the i.n. injection of PBS or IL-3. n=12. **d**, Level of CXCL12 in the BALF of WT or *Cd131*^*−/−*^ mice 24 h after the i.n. injection of PBS or IL-3. n=4-5. **e**, Absolute number of pDCs and neutrophils in the lungs of naive mice 24 h after the i.n. injection of PBS, IgG or anti-CXCL12 in combination with the i.n. injection of PBS (black) or IL-3 (grey). n=8. **f**, Correlation between plasma IL-3 and CXCL12 levels of SARS-CoV-2^+^ patients. n=106. **g**, Level of CXCL12 in the BALF of patients with pulmonary diseases with high or low IL-3 BALF levels. n=13. **h**, Percentage of pDCs in the BALF of patients with pulmonary diseases with high or low CXCL12 BALF levels. n=13. **i**, Percentage of CXCL12^+^ epithelial cells in the lungs of patients with pulmonary inflammation. n=15. **j,** Immunohistochemistry of EpCAM, CXCL12, CD31, and IgG in the lungs of patients with pulmonary inflammation. Data are mean ± s.e.m., *P < 0.05, **P < 0.01, ***P < 0.001, unpaired or Mann Whitney test were used.

I.n. injection of CXCL12 in mice resulted in increased numbers of pDCs but not neutrophils in the lungs (Extended Data Fig. 3c). Additionally, i.n. injection of CXCL12-neutralizing antibodies prevented pDC influx into the lungs of WT mice upon IL-3 injection, whereas no difference was observed for neutrophils (Fig. 2f). In SARS-CoV-2^+^ patients, plasma IL-3 levels strongly correlated with plasma CXCL12 levels, but not with plasma IL-6, TNF and CRP levels (Fig. 2g and Extended Data Fig. 4a-c). CXCL12 plasma levels were also not correlated with circulating pDC number (Extended Data Fig. 4d). In BALF of patients with different respiratory diseases, IL-3 positively correlated with CXCL12 levels (Fig. 2h) and high levels of CXCL12 were associated with increased percentages of pDCs (Fig. 2i).

We found that only CD45^−^ non-haematopoietic cells expressed the α-chain of the IL-3 receptor (CD123) in the lungs of naive mice (Extended Data Fig. 5a). Likewise, only CD45^−^ cells secreted CXCL12 after IL-3 stimulation (Extended Data Fig. 5b). Flow cytometry analyses revealed that only epithelial cells were found to overexpress CXCL12 in the lungs of mice upon *ex vivo* IL-3 stimulation (Extended Data Fig. 5c). As well, CXCL12 was only expressed by CD326^+^ CD123^+^ epithelial cells in human lungs (Fig 2j-k and Extended Data Table 5).

Collectively, our study revealed that plasma IL-3 levels might allow risk stratification in patients with SARS-CoV-2 infections. We therefore propose IL-3 as a predictive marker for disease severity and clinical outcome. Based on its ability to improve local antiviral defence mechanisms by recruiting pDCs, recombinant IL-3, or CD123 receptor agonists, may therefore have the potential as novel therapeutic agents in SARS-CoV-2 infected patients.

## Supplemental Information

### Materials & Methods

#### Animals

Balb/c (WT), C57Bl/6J (WT) (Janvier, Le Genest-Saint-Isle, France) and *Cd131*^*−/−*^ (C57Bl/6J background, bred in-house) mice were used in this study. Majority of the mice were 8-12 weeks old when sacrificed. All animal protocols were approved by the animal review committee from the university hospital Dresden and Erlangen and the local governmental animal committee.

#### Mouse infection

Naive mice were anesthetized with isoflurane and infected intra-nasally with 8μg of CpG (Enzo Life Sciences, Farmingdale, NY, USA), 400ng of recombinant IL-3 (R&D Systems, Minneapolis, MN, USA), 500ng of recombinant CXCL12 (Peprotech, Rocky Hill, NJ, USA), 50μg of IgG or anti-CXCL12 (R&D systems). For the pDCs depletion experiment, 150μg of IgG or anti-CD317 (Miltenyi) were injected intravenously 15h hour before IL-3 and CpG injection.

#### Murine leukocytes isolation

After lungs harvest, single cell suspensions were obtained as follows: perfused lungs were cut in small pieces and subjected to enzymatic digestion with 450 U/ml collagenase I (Sigma Aldrich), 125 U/ml collagenese IX (Sigma Adrich), 60 U/ml hyaluronidase (Sigma Aldrich), 60 U/ml Dnase (Sigma Aldrich) and 20 mM Hepes (Thermo Fisher Scientific, Waltham, MA, USA) for 1 hour at 37°C while shaking. Broncho-alveolar lavage (BAL) was performed by flushing the lungs with 2 × 1 ml of PBS to retrieve the infiltrated and resident leukocytes. Total viable cell numbers were obtained using Trypan Blue (Carl Roth).

#### Quantitative RT-PCR

Real-time PCR was performed as previously described^1^. Briefly, RNA was extracted from whole tissue by RNeasy mini kit (Qiagen, Venlo, Netherlands). Complementary DNA was reverse transcribed from 1 μg total RNA with Moloney murine leukemia virus reverse transcriptase (Thermo Fisher Scientific) using random hexamer oligonucleotides for priming (Thermo Fisher Scientific). The amplification was performed with a Biorad CFX-Connect Real-time-System (Thermo Fisher Scientific) using the SYBR Green (Eurogentec, Seraing, Belgium) or TaqMan (Thermo Fisher Scientific) detection system. Data were analyzed using the software supplied with the Sequence Detector (Life Technologies). The mRNA content for *Cxcl12*, *Ccl2*, *Cxcl9* and *Ccl21* was normalized to the hypoxanthine-guanine phosphoribosyltransferase (*Hprt*) mRNA for mouse genes. Gene expression was quantified using the ∆∆Ct method. The expression level was arbitrarily set to 1 for one sample from the PBS group, and the values for the other samples were calculated relatively to this reference. The sequence of primers is in Extended Data Table 3.

#### Cytokine detection

<u>Mouse:</u> Secreted CXCL12 (R&D systems), IFNλ (R&D systems) and IFNα (R&D systems) were measured by ELISA according to the manufacturer’s instructions. <u>Human:</u> Secreted CXCL12 (R&D Systems), IL-3 (R&D Systems), IL-6 (Biolegend) and TNF (Biolegend) were measured by ELISA according to the manufacturer’s instructions.

#### Lung cells stimulation *in vitro*

Lung cell suspensions from naive mice were cultured in RPMI-1640 GlutaMax supplemented with 10% FCS, 25mM of Hepes, 1 mM sodium pyruvate, 100U/ml of Penicillin-Streptomycin and 20μg/ml of Gentamicin at 37°C in the presence of 5% CO2. Lung cell suspensions were stimulated in 12-well plates (10^6^ cells/ml) during 24h by IL-3 (20ng/ml). Then supernatants were collected for cytokine measurement. CD45^−^ and CD45^+^ cells were purified from lungs of naive mice using CD45 microbeads (Miltenyi Biotec), according to the manufacturer’s instructions.

#### Immunohistochemistry

For CXCL12 and EpCAM permanent immunohistochemistry, the staining was performed as previously described^2^. The characteristics of the respective patients are detailed in Extended Data Table 2. In brief, formalin-fixed, paraffin-embedded lung tissues were deparaffinized by xylene two times for 15 min. The tissue was rehydrated using decreasing concentrations of ethanol (100%, 96%, 85% and 70%) for 2 min each. Antigen retrieval was performed using Target Retrieval Solution (DakoCytomation) at pH 9.0 at 95°C for 20 min followed by cooling for 20 min at RT. As a washing buffer between the incubation steps, 1xTBS pH 7.6 was used. The slides were blocked by hydrogen peroxide (7.5%, Sigma-Aldrich), avidin-biotin-block (Vector Laboratories, Burlingame, CA, USA) and 10% donkey normal serum (DNS, Vector Laboratories) in TBS for 10 min. The primary antibodies diluted in 2.5% DNS (rabbit anti-human EpCAM cat no. ab71916, Abcam, 1:300; mouse anti-human CXCL12 cat. no. MAB350, R&D Systems, 1:150) and isotype control antibodies in corresponding concentrations were detected using either the RTU Vectastain Elite ABC Kit anti-mouse/rabbit (for EpCAM and CXCL12; Vector Laboratories) and NovaRed substrate (Vector Laboratories) as a substrate. The slides were counterstained with Gill-III hematoxylin (Merck), dehydrated and mounted with VectaMount permanent mounting medium (Vector Laboratories). The sections were analyzed using a DM6000 B microscope (Leica).

#### Flow Cytometry

The following antibodies were used for flow cytometric analyses: <u>Mouse:</u> anti-CD317-BV650 (Biolegend), anti-Ly6C-FITC (BD Biosciences), anti-B220-BUV737 (BD Biosciences), anti-CD11c-PerCP Cy5.5 (Biolegend), anti-CD11b-PE CF594 (BD Biosciences), anti-F4/80-BV510 (BD Biosciences), anti-Ly6G-BUV395 (BD Biosciences), anti-SiglecH-Pacific Blue (BD Biosciences), anti-CD45.2-BV786 (BD Biosciences), anti-MHCII-BV711 (BD Biosciences), anti-CD19-BV421 (BD Biosciences), anti-CD3-PerCP Cy5.5 (Biolegend), IgG2a-PE (Biolegend), anti-CD146-FITC (Biolegend), anti-CD31-Pacific Blue (Biolegend), anti-CD326-PE-CF594 (Biolegend), anti-CD326-BV650 (BD Biosciences), anti-CD123-PE (Biolegend). <u>Human:</u> anti-CD45-Pacific Blue (Biolegend), anti-CD45-BV786 (BD Biosciences), anti-CD303-PerCP Cy5.5 (Biolegend), anti-HLA-DR-BUV395 (BD Biosciences), anti-CD11c-BV711 (BD Biosciences), anti-CD326-PercCP Cy5.5 (Biolegend), anti-CD31-BV711 (BD Biosciences), anti-CD14-BUV737 (BD Biosciences), anti-CD16-BV421 (BD Biosciences), anti-CD11b-BV711 (BD Biosciences), anti-CD15-PE (BD Biosciences). Anti-CXC12-PE (R&D Systems) and IgG1-PE (R&D Systems) were used for mouse and human. Staining for intracellular cytokines was performed using BD Cytofix/Cytoperm Plus Kit (BD Biosciences). Data were acquired on a Celesta (BD Biosciences) flow cytometer and analyzed with FlowJo 10 (FlowJo LLC, Ashland, OR, USA).

#### Human specimen

This prospective, multicentric, observational clinical study was first approved by the local ethics committee on February, 1^st^ 2016 (UKER 10_16 B), and modified on April, 28^th^ 2020 (UKER 174_20 B). Departments from 3 additional University Hospitals in Germany participated in this study. Prospective measurements of Interleukin-3 and analysis of patients participating in the trial have been conducted between April 1^st^ 2020 and June 4^th^ 2020. The observational clinical studies were conducted in the medical wards and intensive care units of the (i) University Hospital of Erlangen, Germany; (ii) University Hospital of Essen, Germany; (iii) University Hospital of Giessen, Germany; and (iv) University Hospital of Bonn, Germany. Study patients or their legal designees signed written informed consent. In total, 106 (32 non-severe; 32 severe; 42 recovered) patients positive for SARS-CoV-2 PCR from oral swabs, oral fluid, or bronchoalveolar lavage fluid were enrolled in this trial. Blood samples were collected at the onset of symptoms (≤ 24 hours), and 1, 2, 3, 4, 5, 6, or 7 days later; or after recovery from SARS-CoV-2 infection. Patients with high viral load are patients with CT levels above the median of all patients (Median CT levels: 32.63). 20 healthy donors served as controls. *Blood:* After blood collection, plasma of all study participants was immediately obtained by centrifugation, transferred into cryotubes, and stored at −80°C until further processing. For flow cytometry analysis, red blood cells were removed by centrifuging blood cells 7 min at 400g without break in Leukosep tube (Greiner, Kremsmünster, Austria). Leukocytes were then stained 20 min at 4°C in dark and fixed 1 hour with BD Cytofix buffer (BD Biosciences). *Bronchoalveolar-lavage fluid specimen (13 patients):* After written informed consent and in agreement with the local ethics review board of the University of Erlangen (UKER no. 4147) the segmental bronchi of patients scheduled for fiber-optic bronchoscopy due to various inflammatory and non-inflammatory conditions were flushed with 100ml sterile 0.9% saline fluid. Fluid has been obtained and processed for flow cytometric analysis of leukocyte surface markers. In addition, after centrifugation the supernatants have been stored at −80°C until further processing. Patients with low IL-3 or low CXCL12 are patients with a level of IL-3 or CXCL12 under the mean of all the patients. Patients with high IL-3 or high CXCL12 are patients with a level of IL-3 or CXCL12 above the mean of all the patients. *Lung tissue specimen (15 patients):* The study was performed at the University of Erlangen in Germany. Patients were selected within the framework of the thoracic surgery board. The patients who underwent surgery and gave their approval were included in this study. The study was performed in agreement with the local ethics review board of the University of Erlangen (UKER 10_16 B, UKER 339_15 Bc; UKER 56_12B; DRKS-ID: DRKS00005376). Patients’ confidentiality was maintained. The surgery consisted of a wedge resection of the lung or lobectomy. Subsequently, tissue samples were taken from the surgically removed material and transported into the laboratory under standardized conditions (at 4°C, in Ringer’s solution) for further preparation and analysis. Samples were taken from the non-pathological area from the lung for further analysis. For flow cytometry analysis, separate lung tissue sections were cut into small pieces and subjected to enzymatic digestion with 450 U/ml collagenase I, 125 U/ml collagenase XI, 60 U/ml DNase I and 60 U/ml hyaluronidase (Sigma-Aldrich, St. Louis, MO) for 1 h at 37°C while shaking at 750 rpm after which they have been homogenized through a 40μm nylon mesh for flow cytometric analysis. Total viable cell numbers were obtained using Trypan Blue (Cellgro, Mediatech, Inc, VA).

#### Statistics

Results were expressed as mean ± S.E.M. and expressed as identified in legends. For comparing 2 groups, statistical tests included unpaired, 2-tailed parametric t tests with Welch’s correction (when Gaussian distribution was assumed), unpaired, 2-tailed nonparametric Mann-Whitney tests (when Gaussian distribution was not assumed) or paired, 2-tailed parametric t tests. P values of 0.05 or less were considered to denote significance.

## Extended Data Figures

**Extended Data Fig. 1.**
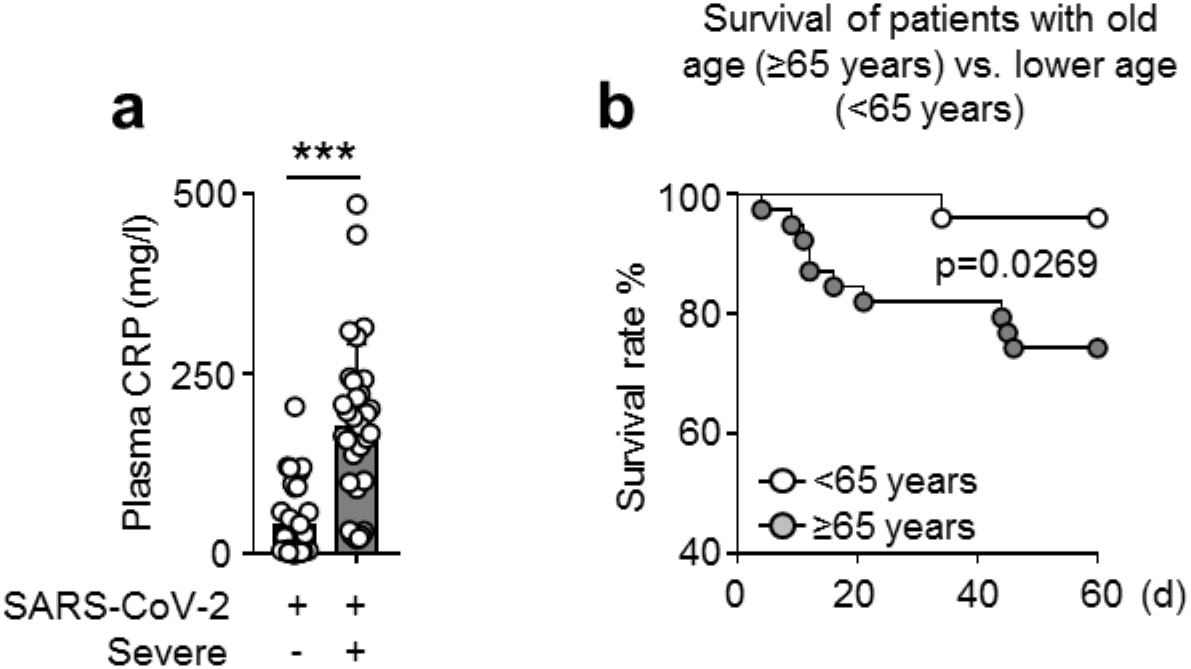
Plasma CRP levels and survival of SARS-CoV-2^+^ patients with old (≥65 years) vs. lower (<65 years) age. **a**, Plasma CRP levels in SARS-CoV-2^+^ patients with severe or non-severe disease. Mann Whitney test. n=64. **b**, Kaplan-Meier analysis showing the survival of SARS-CoV-2^+^ patients with old (≥65 years) or younger (<65 years) age. n=64. Data are mean ± s.e.m., *P < 0.05, **P < 0.01, ***P < 0.001.

**Extended Data Fig. 2.**
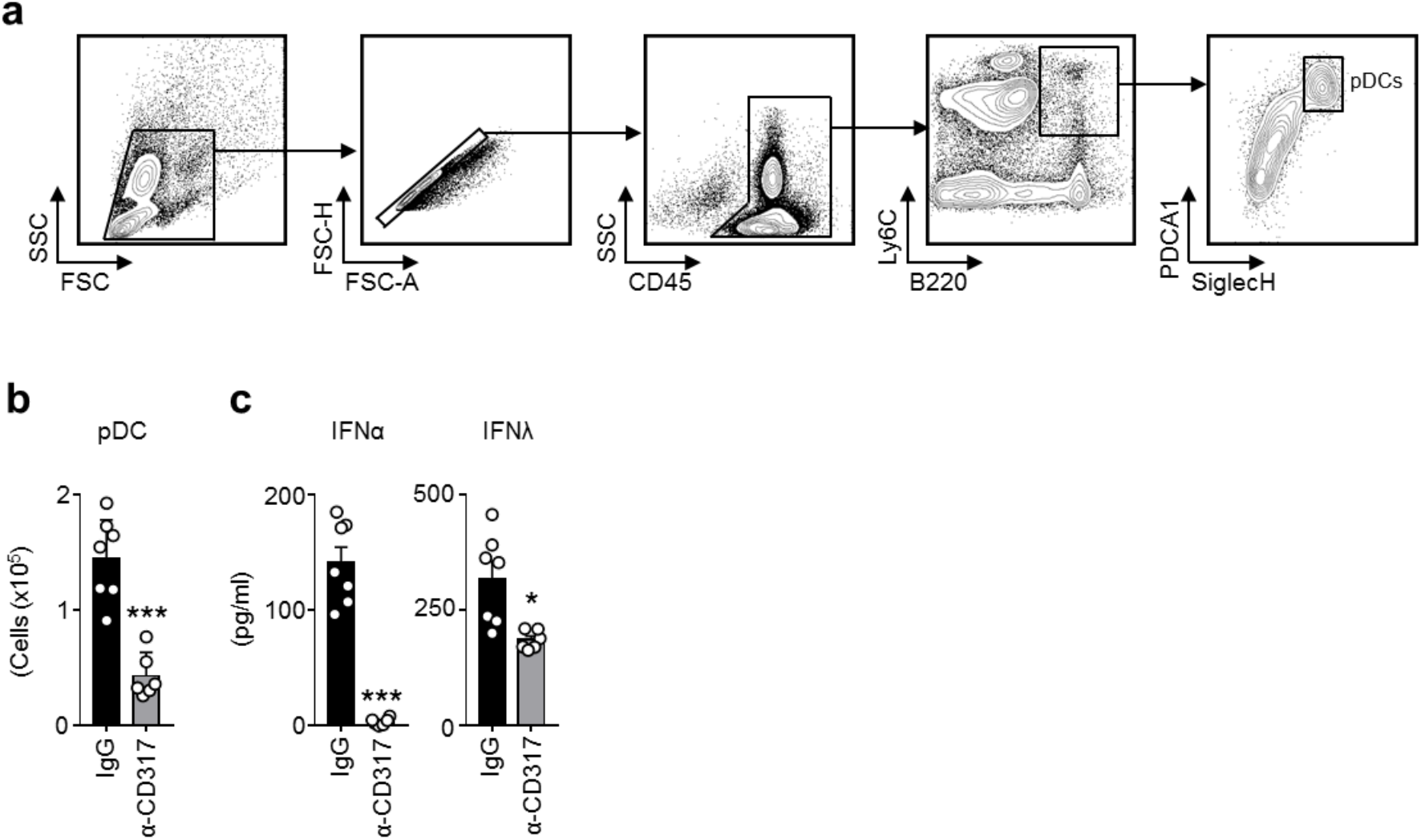
Depletion of pDCs in mice pre-treated with IL-3 induces reduced BALF IFNα and IFNλ levels upon i.n. CpG administration. **a**, Gating strategy used for pDCs. **b-c**, Absolute numbers of pDCs in the lungs (**b**) and levels of IFNα and IFNλ in the BALF (**c**) of naive mice intravenously injected with IgG or anti-CD317 15 h before the injection of IL-3 and CpG. n=7. Data are mean ± s.e.m., *P < 0.05, **P < 0.01, ***P < 0.001, unpaired, 2-tailed Student’s t test using Welch’s correction for unequal variances was used.

**Extended Data Fig. 3.**
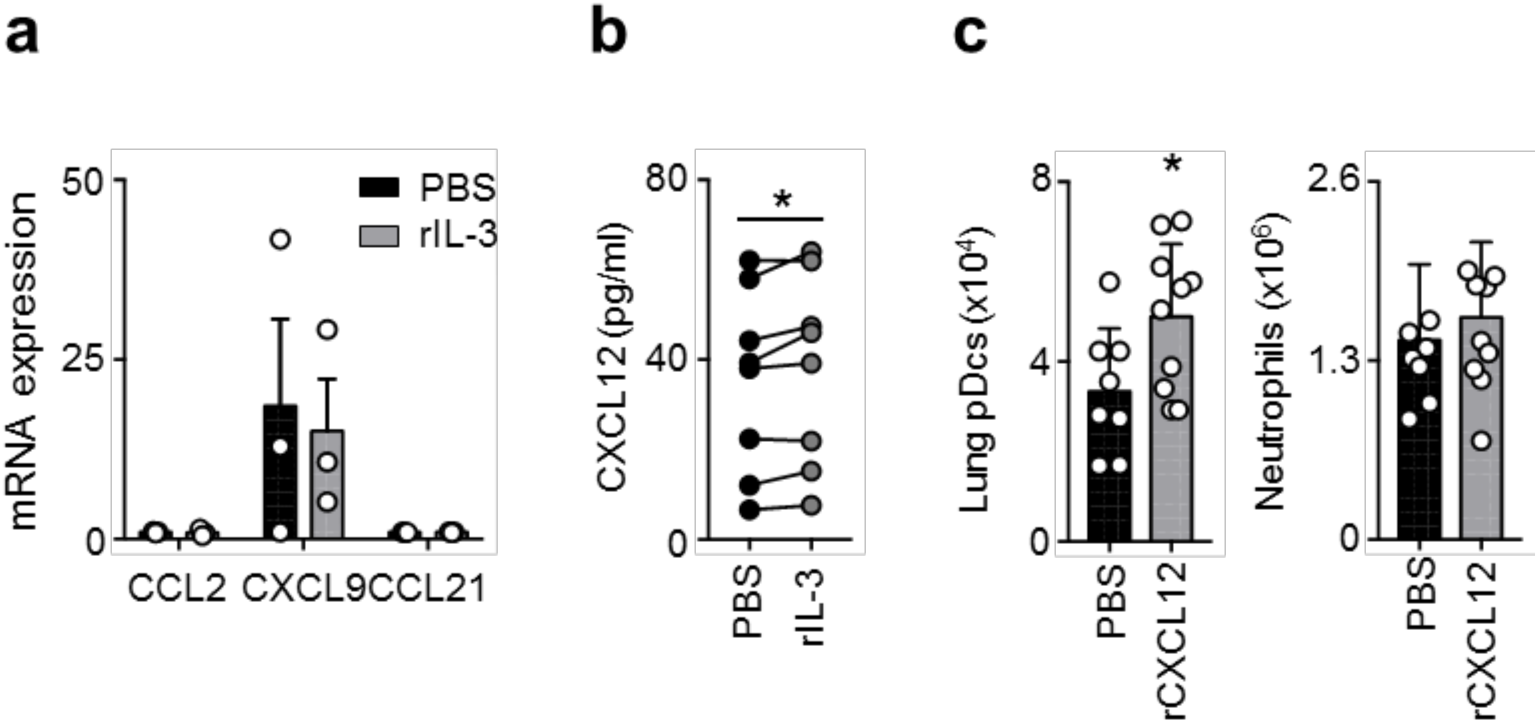
Interleukin-3 induces CXCL12. **a**, Relative mRNA expression of *Ccl2*, *Cxcl9* and *Ccl21* in the lungs of naive mice 24 h after the i.n. injection of PBS or IL-3. n=3. **b**, Levels of CXCL12 in the supernatant of lungs cells from naive mice 24 h after *ex vivo* stimulation with or without IL-3. n=8-14. **c**, Absolute numbers of pDCs or neutrophils in the lungs of naive mice 24 h after the i.n. injection of PBS or CXCL12. n=7-10. Data are mean ± s.e.m., *P < 0.05, **P < 0.01, ***P < 0.001, paired, 2-tailed Student’s t test and unpaired, 2-tailed Student’s t test using Welch’s correction for unequal variances was used.

**Extended Data Fig. 4.**
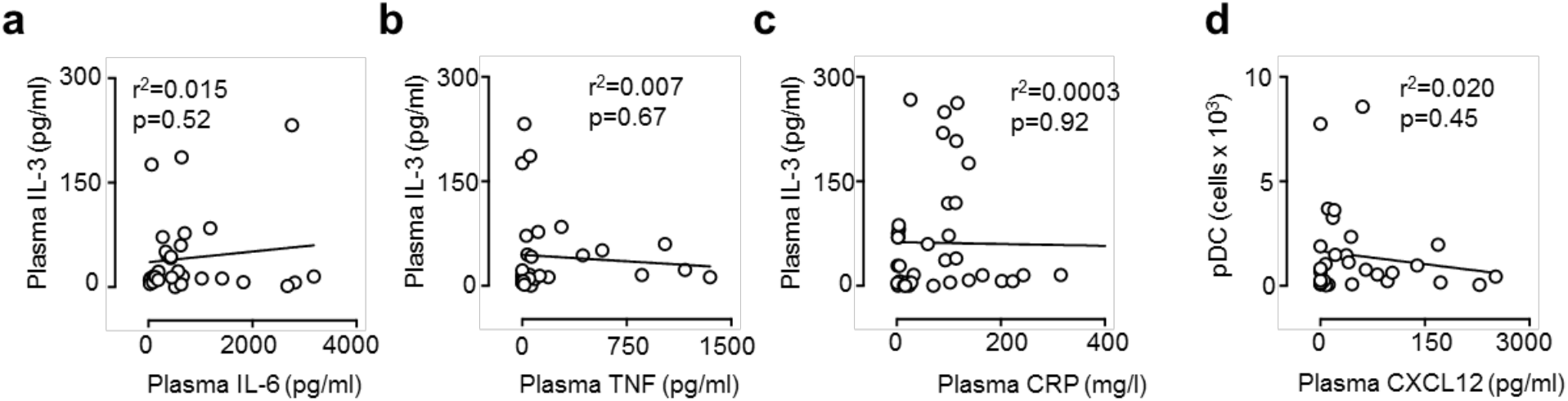
Plasma IL-3 levels do not correlate with plasma IL-6, TNF, CRP levels and circulating pDCs. **a-c,** Correlation between plasma IL-3 levels and plasma IL-6 (**a**), TNF (**b**) and CRP (**c**) levels in SARS-CoV-2^+^ patients. n=32. **d**, Correlation between plasma CXCL12 levels and the amount of circulating pDCs in SARS-CoV-2^+^ patients. n=32.

**Extended Data Fig. 5.**
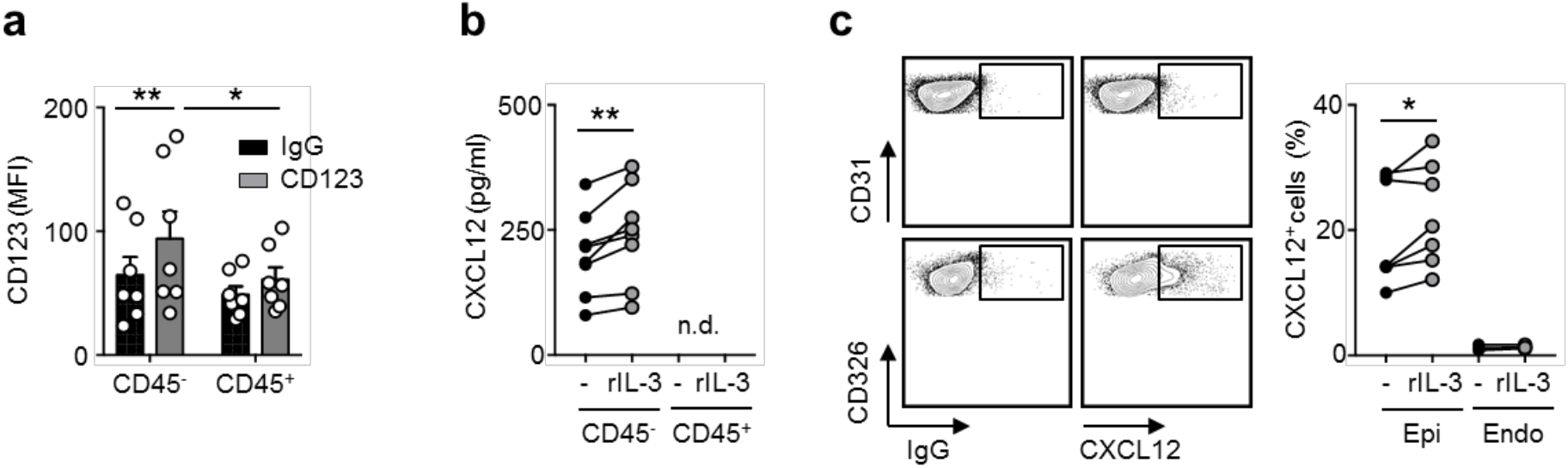
Interleukin-3 induces CXCL12 in lung epithelial cells. **a**, CD123 expression (MFI) on the surface of CD45^−^ and CD45^+^ cells from the lungs of naive mice. n=8. **b**, Level of CXCL12 in the supernatant of CD45^−^ or CD45^+^ cells purified from the lungs of naive mice 24 h after *ex vivo* stimulation with or without IL-3. n=8. **c**, Representative dot plot (left) and percentage (right) of CXCL12^+^ epithelial or endothelial cells in the lungs of mice 24 h after *ex vivo* stimulation with or without IL-3. n=7. Data are mean ± s.e.m., *P < 0.05, **P < 0.01, ***P < 0.001, paired, 2-tailed Student’s t test and unpaired, 2-tailed Student’s t test using Welch’s correction for unequal variances was used.

**Extended Data Table 1:**
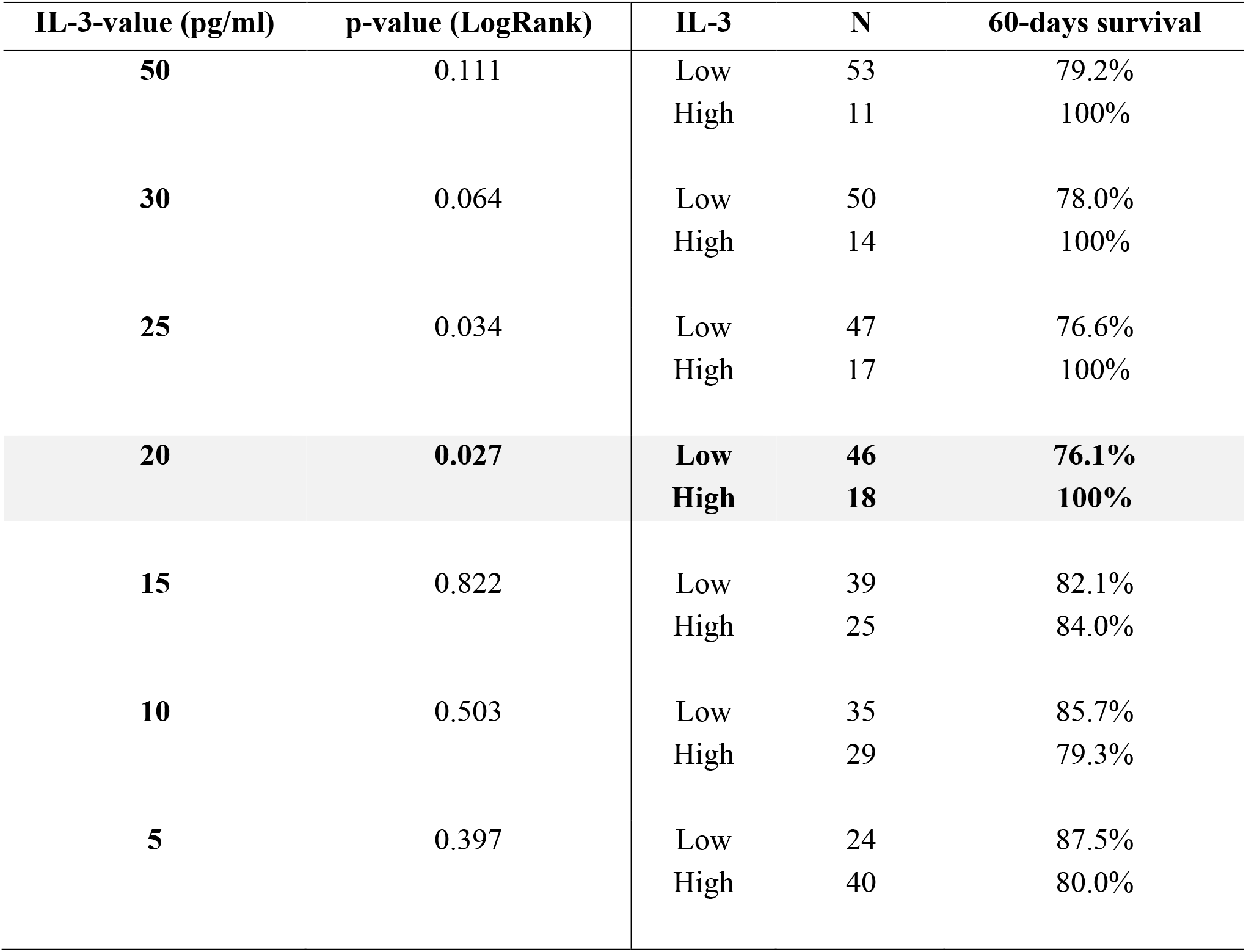
Minimum p-value approach for IL-3 (n=64).

| IL-3-value (pg/ml) | p-value (LogRank) | IL-3 | N | 60-days survival |
| --- | --- | --- | --- | --- |
| **50** | 0.111 | Low | 53 | 79.2% |
|  |  | High | 11 | 100% |
| **30** | 0.064 | Low | 50 | 78.0% |
|  |  | High | 14 | 100% |
| **25** | 0.034 | Low | 47 | 76.6% |
|  |  | High | 17 | 100% |
| **20** | **0.027** | **Low** | **46** | **76.1%** |
|  |  | **High** | **18** | **100%** |
| **15** | 0.822 | Low | 39 | 82.1% |
|  |  | High | 25 | 84.0% |
| **10** | 0.503 | Low | 35 | 85.7% |
|  |  | High | 29 | 79.3% |
| **5** | 0.397 | Low | 24 | 87.5% |
|  |  | High | 40 | 80.0% |

**Extended Data Table 2:**
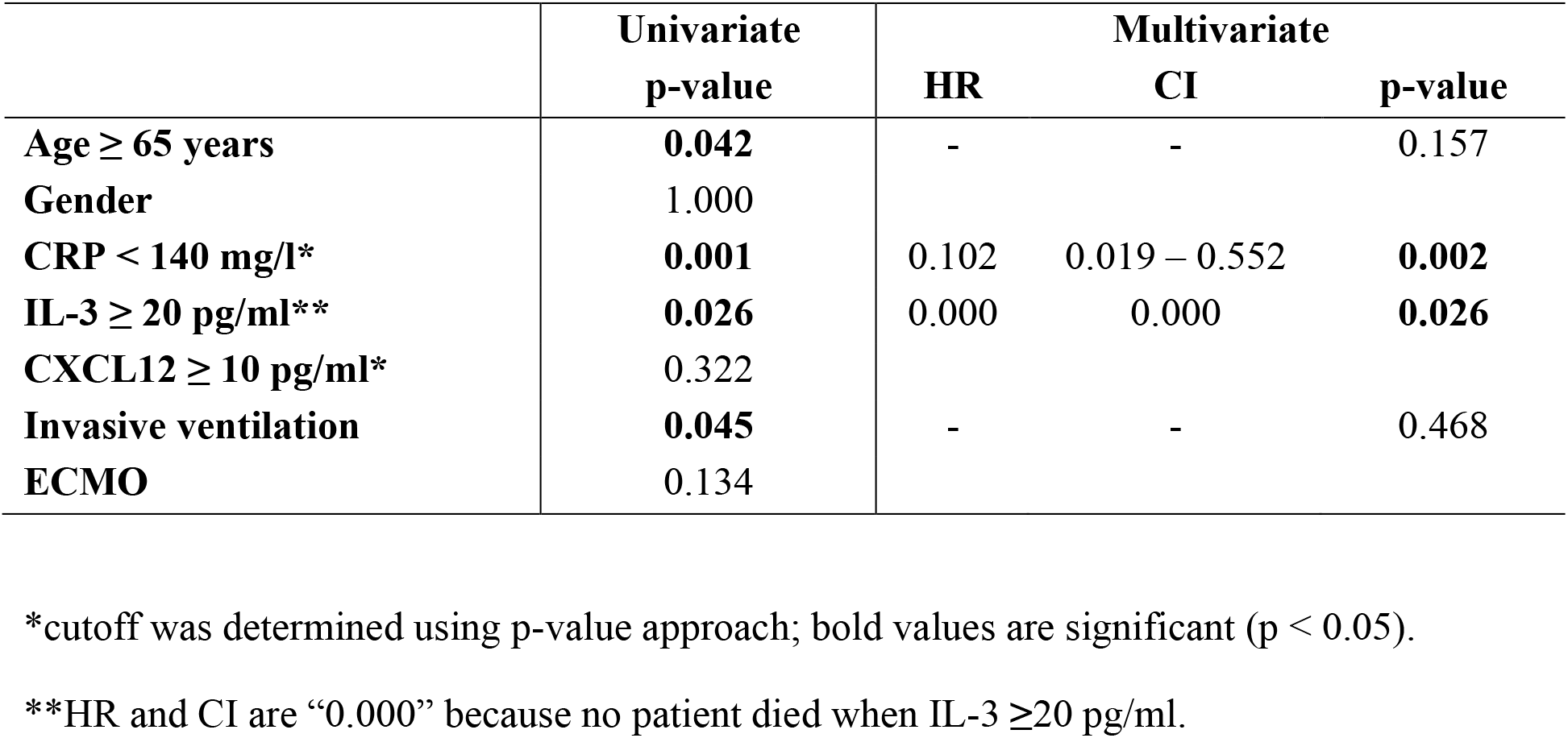
Multivariate analysis of the impact of different risk factors on mortality following SARS-CoV-2-infection (n=64).

**Extended Data Table 3:**
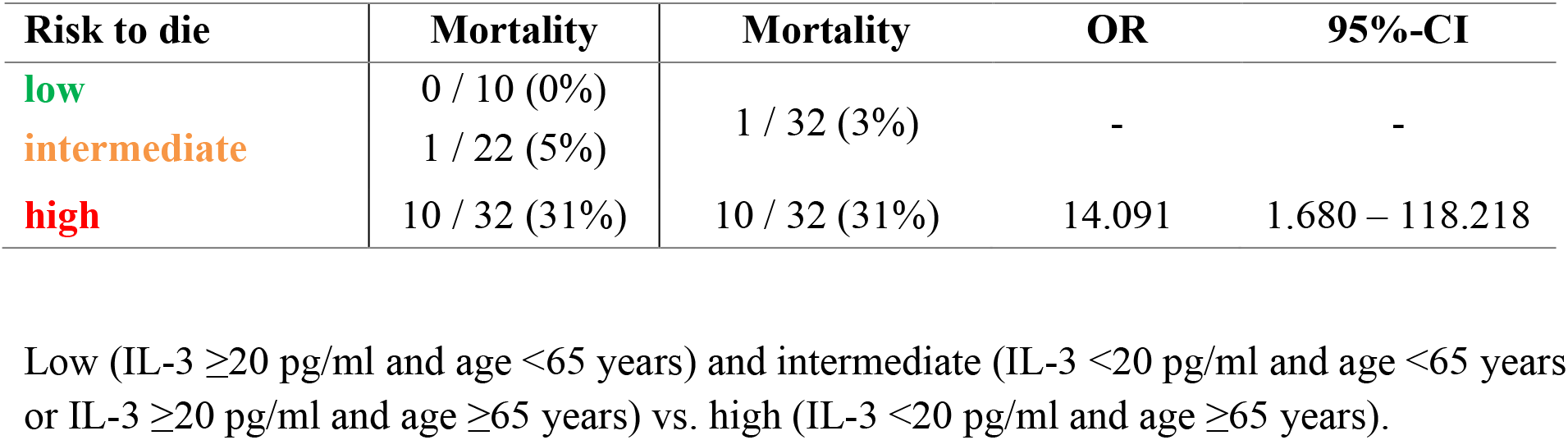
Risk to die from SARS-CoV-2 according to risk groups.

| Risk to die | Mortality | Mortality | OR | 95%-CI |
| --- | --- | --- | --- | --- |
| low | 0 / 10 (0%) | 1 / 32 (3%) | - | - |
| intermediate | 1 / 22 (5%) |  |  |  |
| high | 10 / 32 (31%) | 10 / 32 (31%) | 14.091 | 1.680 – 118.218 |

Low (IL-3 $\geq 20$ pg/ml and age <65 years) and intermediate (IL-3 <20 pg/ml and age <65 years or IL-3 $\geq 20$ pg/ml and age $\geq 65$ years) vs. high (IL-3 <20 pg/ml and age $\geq 65$ years).

**Extended Data Table 4:**
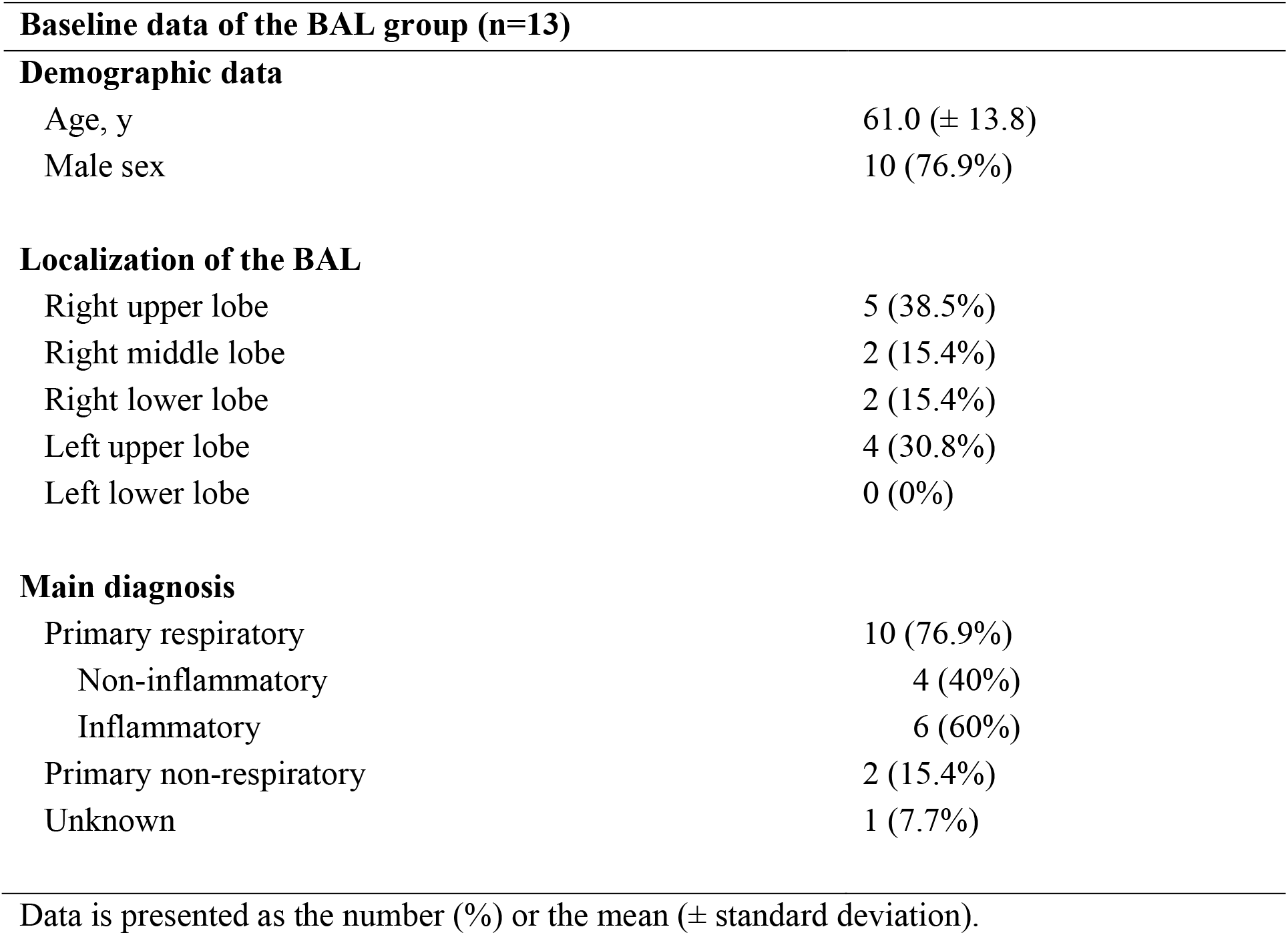
Baseline data of patients scheduled for elective bronchoalveolar lavage.

**Demographic data**
|  |  |
| --- | --- |
| Age, y | 61.0 (± 13.8) |
| Male sex | 10 (76.9%) |
| **Localization of the BAL** |  |
| Right upper lobe | 5 (38.5%) |
| Right middle lobe | 2 (15.4%) |
| Right lower lobe | 2 (15.4%) |
| Left upper lobe | 4 (30.8%) |
| Left lower lobe | 0 (0%) |
| **Main diagnosis** |  |
| Primary respiratory | 10 (76.9%) |
| Non-inflammatory | 4 (40%) |
| Inflammatory | 6 (60%) |
| Primary non-respiratory | 2 (15.4%) |
| Unknown | 1 (7.7%) |

Data is presented as the number (%) or the mean ( ± standard deviation).

**Extended Data Table 5:**
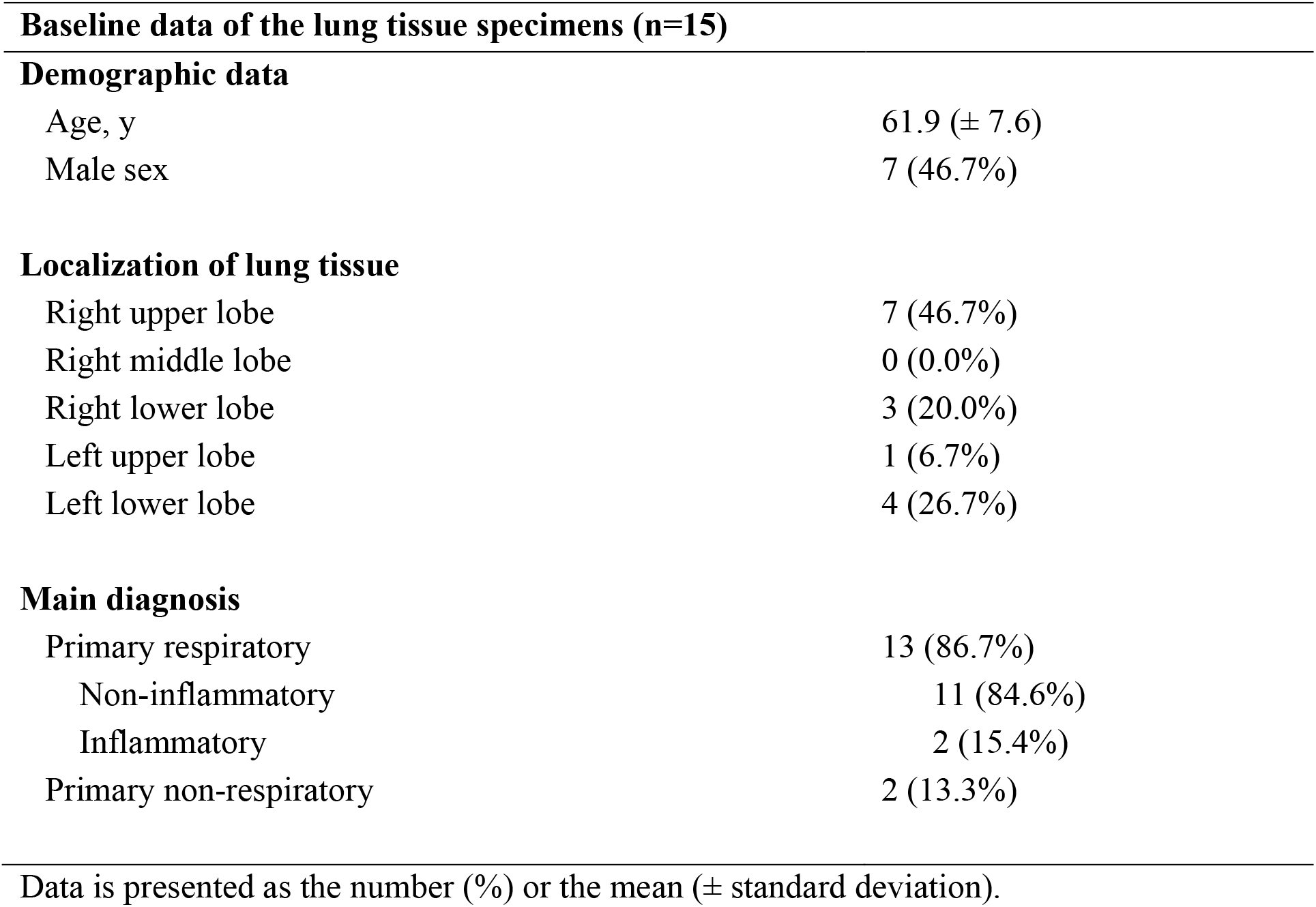
Baseline data of patients scheduled for elective thoracic surgery to obtain lung tissue.

| **Baseline data of the lung tissue specimens (n=15)** |  |  |
| --- | --- | --- |
| **Demographic data** |  |  |
| Age, y | | 61.9 (± 7.6) |
| Male sex |  | 7 (46.7%) |
| **Localization of lung tissue** |  |  |
| Right upper lobe |  | 7 (46.7%) |
| Right middle lobe |  | 0 (0.0%) |
| Right lower lobe |  | 3 (20.0%) |
| Left upper lobe |  | 1 (6.7%) |
| Left lower lobe |  | 4 (26.7%) |
| **Main diagnosis** |  |  |
| Primary respiratory |  | 13 (86.7%) |
| Non-inflammatory |  | 11 (84.6%) |
| Inflammatory |  | 2 (15.4%) |
| Primary non-respiratory |  | 2 (13.3%) |

Data is presented as the number (%) or the mean ( ± standard deviation).

**Extended Data Table 6:**
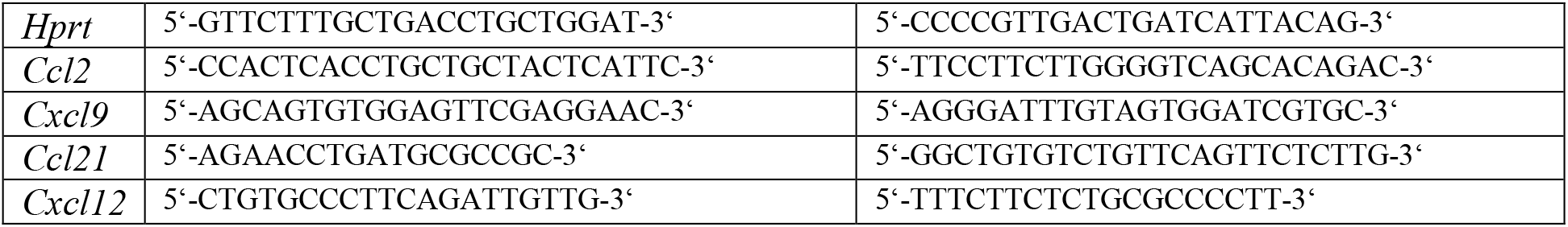
Mouse primer.

